# Systems modeling of mitochondrial dynamics in different exercise regimes

**DOI:** 10.1101/2025.10.27.684912

**Authors:** Ali Khalilimeybodi, Lingxia Qiao, Allen Leung, Andrew D. McCulloch, Simon Schenk, Padmini Rangamani

## Abstract

Exercise stimulates skeletal muscle signaling and mitochondrial metabolism. Emerging evidence shows that mitochondrial dynamics (i.e., fission and fusion) could be regulated by exercise. Yet, there are knowledge gaps on the following questions: (i) which upstream signals are necessary and sufficient to bias mitochondria toward fission versus fusion? (ii) How does cellular energy status and ROS partition control between DRP1 and MFN/OPA1? And (iii) which combinations of intensity and duration produce similar cytosolic signals but distinct mitochondrial remodeling? To address these gaps, we developed an integrative computational framework that connects exercise regimens to mitochondria fission-fusion machinery by linking blood-myofiber energetics in cytosol and mitochondria to skeletal muscle signaling network. The influence of three exercise regimen (i.e., sprint, resistance, and endurance) on mitochondrial fission and fusion was simulated. Classified qualitative validation of signaling network model against studies not used in developing the model achieved 80% accuracy. The model predicts regimen-specific dynamics starting with acute DRP1-driven fission during exercise followed by MFN1/2–OPA1-mediated re-fusion as energy stress declines, consistent with a cyclical triage-then-rebuild paradigm. Changes are most pronounced and sustained with endurance, sharp but brief with sprint, and minimal with resistance. Global sensitivity analysis identified AMPK/PGC-1α→MFN1/2 as dominant fusion drivers, ROS and AMPK→MFF/DRP1 as primary fission switches, and Ca²⁺-calmodulin, ERK, and LKB1/AMPK as shared regulators of fission and fusion. Our model also predicts that an endurance base, augmented with 1–2 weekly high intensity interval traning (HIIT)/ sprint interval training (SIT) sessions could maximize AMPK-ROS pulses and mitochondrial fission-fusion. This framework unifies muscle’s signaling logic with the energetic state to explain how intensity-volume combinations, bout spacing, and kinase modulation tune mitochondrial remodeling, yielding testable predictions for optimizing training and adjuvant therapies for enhanced human performance.

## Introduction

Exercise is a potent physiological perturbation to skeletal muscles, eliciting coordinated shifts in signaling cascades, metabolic flux, gene transcription, and cellular phenotype ^1,2^. Across different types of exercise modalities including resistance, endurance, and sprint, acute bouts of physical activity remodel the phosphoproteome including the regulatory kinases AMPK, mTOR, and PKA ^3^. Yet, superimposed on this common core of change in phosphorylation patterns are modality-specific patterns, particularly evident during recovery ^1–3^. ATP generation in muscles is matched to contractile demand through substrate-level (anaerobic) and oxidative (aerobic) phosphorylation ^4^. The relative engagement of ATP-generating pathways is dictated by the exercise task. During brief, all-out efforts lasting only seconds (e.g., sprinting), ATP generation arises mainly from phosphocreatine breakdown and glycogen-to-lactate flux ^5,6^, reflecting substrate-level (anaerobic) phosphorylation. In contrast, during exercise sustained for minutes to hours, ATP is supplied predominantly through oxidative metabolism of carbohydrate and fat ^7,8^, characteristic of the aerobic energy system. Transcriptomic programs likewise diverge by exercise modality. For example, sprint exercise preferentially upregulates stress-response, muscle-adaptation, and immune-related signatures ^9^, while resistance and endurance training enhance genes linked to stress response, kinase activity, and metabolic processes ^10^. This systematic view of exercise which links upstream signals and energy demand to mitochondrial network remodeling in each exercise modality motivates the modeling framework developed in this work.

Mitochondria sit at the center of the exercise response, coordinating ATP supply, redox signaling, and structural remodeling in muscles. During sustained, high-intensity efforts, mitochondria supply the majority of ATP via oxidative phosphorylation, placing them as principal mediators of performance and recovery dynamics ^4^. Classical training studies demonstrate increases in mitochondrial enzymes and oxygen uptake capacity with repeated endurance sessions ^11,12^. Granata et al. showed that manipulating training variables (volume, intensity) yields distinct effects on mitochondrial content versus function ^13^. An acute session of high intensity interval training (HIIT) elevates nuclear PGC-1α and initiates a mitochondrial gene program in human skeletal muscle ^14^. This is in line with rapid exercise-induced surges in PGC-1α transcripts and protein ^15,16^, which can produce robust mitochondrial remodeling after repeated sessions. Mitochondrial dynamics, particularly fusion and fission, are integral to these adaptations and are differentially engaged by exercise dose and timing ^17^. Muscle contractions activate AMPK, cAMP/PKA, and p38 MAPK pathways that converge on PGC-1α; downstream and PGC-1α increases expression of fusion machinery MFN1/MFN2 and OPA1 ^18–21^. In parallel, fission is mediated by the large GTPase DRP1 ^22^. Acute endurance exercise increases DRP1 phosphorylation, with the signal subsiding during recovery. This temporal pattern is consistent with early network fragmentation to segregate damaged segments, followed by re-fusion and biogenesis ^23^. AMPK and Akt can promote DRP1 activity ^24^, whereas PKA phosphorylation inhibits DRP1 and restrains fission ^25^. Resistance exercise presents a different picture: hypertrophy can dilute volume density, yet multiple studies indicate improved mitochondrial function even when classical content markers do not rise. For example, low-load, blood-flow-restricted and high-load resistance training both increase mitochondrial protein synthesis and respiration with no change in citrate synthase activity, implying functional enhancement independent of bulk content ^26^. Together, these data position mitochondria as both the means (ATP supply) and a target (biogenesis, dynamics, mitophagy) of training adaptation, with distinct exercise modalities differentially weighting these levers.

Systems biology models offer a rigorous way to connect exercise dose (intensity, duration, and modality) with skeletal muscle multiscale physiology, by integrating fast metabolic transients, slower signaling programs, and cumulative structural remodeling in the muscle. Early mechanistic work showed that simple control laws can capture high-resolution bioenergetic data. For example, a linear respiration model that explains the mono-exponential phosphocreatine (PCr) dynamics observed in exercising muscle establishes how mitochondrial oxidative flux tracks ATP demand ^27^. Building on this foundation, oxygen transport-metabolism models linked convective-diffusive O₂ delivery to intracellular ATP/PCr kinetics, shedding light on the conditions in which increasing O₂ supply (e.g., hyperoxia) does or does not accelerate oxidative responses during muscle contractions ^28^. Systems bioenergetic models then unified cellular metabolism with training history, predicting how loading or unloading reprograms enzyme capacities and fuel selection during subsequent exercise ^29^. Parameterized against human biopsy and *in vivo* data, *in silico* studies demonstrated how the malate–aspartate shuttle and initial glycogen stores set the time course of glycolytic vs. oxidative ATP provision during work transitions ^30^. Fiber-type–resolved models further dissected Type I/II recruitment and metabolite transport to reproduce heterogeneity in cytosolic and mitochondrial fluxes under moderate-intensity protocols ^31^. On the signaling side, dynamic models have quantified kinase control, showing that ADP, rather than AMP, dominantly governs AMPK activity during exercise, a key insight for linking energy charge to downstream transcriptional programs ^32^. Recently, a signaling network model explored muscle acute and phenotypic responses to endurance vs. resistance exercise and highlighted mTOR– Akt, AMPK and ROS–NFκB as key signaling mediators ^33^. Modular whole-muscle frameworks have coupled metabolism, kinase cascades, and gene expression to recapitulate acute-to-chronic adaptations following varied exercise stimuli ^34^. In parallel, quantitative models of mitochondrial fission–fusion have mapped how network dynamics govern mitochondrial DNA maintenance and quality control, providing a formal handle on morphology–function coupling ^35^.

In this work, we developed a modular systems biology model that links blood-muscle fiber-organelle exchanges to cytosolic signaling and mitochondrial bioenergetics across different timescales to predict how sprint, resistance, and endurance stimuli regulate mitochondrial dynamics. This model integrates a metabolism module adapted from an established exercise model ^31,34^ and a logic-based ODE signaling module developed from prior knowledge and validated against independent studies. Here, we address three key questions: which upstream signals are necessary and sufficient to bias mitochondria toward fission versus fusion? How do cellular energetic status and ROS partition control between DRP1 and MFN/OPA1? And which combinations of intensity and duration produce distinct mitochondrial remodeling despite similar cytosolic signals? Model simulations generated testable, exercise modality-dependent predictions of metabolites (ATP/ADP, lactate, ROS), kinases (AMPK, mTOR, ERK/p38), and effectors of mitochondrial dynamics (DRP1, MFN1/2, OPA1), allowing us to identify principles governing the fission–fusion balance during different exercise regimes.

## Methods

### Model Overview

The aim of this study was to create a mathematical model that integrates the metabolic and signaling pathways linking exercise modalities to mitochondrial fusion and fission. The metabolic module of the model is adopted from a previously developed mathematical model of cellular metabolic dynamics in skeletal muscle fibers during exercise ^31,34^. The signaling module was developed using a logic-based ordinary differential equation (ODE) approach ^36^ frequently used in modeling large-scale signaling pathways in cardiac ^37–39^ as well as skeletal muscles ^33^. By integrating key mediators that connect skeletal muscle metabolism and signaling, such as ATP/ADP, CPT1, ROS, and GLUT4 into the model, we combined the signaling and metabolic modules to capture mitochondrial dynamics (i.e., fusion and fission) across various exercise regimens. Given our focus on differential dynamics of signaling and metabolic species between exercise modalities, model simulation results are reported as normalized values. The normalized level is obtained by the fractional activation of species divided by the maximum changes in the fractional activation across three different types of exercise regimens.

### Metabolic Module

For metabolic module, we adapted a previously established mechanistic framework that couples cytosolic glycolysis, PCr/Cr buffering, lipid handling, and mitochondrial TCA– OXPHOS to simulate exercise-induced metabolic transients in skeletal muscle ^31^. We analyzed the muscle as a well-mixed tissue compartment without explicit fiber-type differentiation. Protein metabolism was not included in the model, as its contribution to energy production during short-term exercise is minimal ^31^. The model includes distinct compartments for cytosolic and mitochondrial domains, allowing for the simulation of mass transport processes and reaction fluxes that occur during exercise. For the transition from rest to exercise, skeletal muscle was modeled to undergo an activation process capturing enhanced metabolic rate within muscle cells. The metabolic module also accounts for transport processes between the blood and muscle, enabling simulation of how blood-borne substrates and metabolites are utilized by the muscle during exercise ^31^. The transport processes between the blood and muscle fibers were modeled using dynamic mass balance equations. These equations describe the transport of chemical species from the blood to the cytosol and from the cytosol to the mitochondria. Mass transport was influenced by both passive diffusion and facilitated transport. The metabolic reaction fluxes in the cytosol and mitochondria were governed by reversible enzyme kinetics, with the forward and backward reaction rates adjusted according to the experimental data on substrate and product concentrations ^31^. The metabolic model includes a series of reactions modeling the breakdown of carbohydrates and fats, as well as the synthesis of ATP through oxidative phosphorylation in the mitochondria. The parameters related to activation coefficients were optimized by fitting model outputs to experimental data. Model validation was performed by comparing simulated metabolite concentrations with published experimental data on muscle fiber activation patterns during exercise ^31,34^.

#### Minimal Model for reactive oxygen species

To estimate reactive oxygen species (ROS) changes during exercise, we incorporated both mitochondrial and cytosolic ROS generation mechanisms in the model. For mitochondrial ROS, as demonstrated in previous studies ^40^, we used the following function

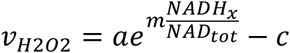

to represent the ROS production rate caused by the NADH/NAD⁺ ratio in skeletal muscle mitochondria. This exponential nonlinear regression function provides a good fit to the experimental training data and can be adjusted or normalized as needed ^40^. During exercise, the PI3K/Akt and PKC pathways, along with mechanical stress, contribute to increased ROS production ^41–43^. The PI3K/Akt pathway increases mitochondrial activity, resulting in greater electron leakage from the electron transport chain and subsequent ROS generation ^44^. PKC activation, driven by elevated calcium levels and diacylglycerol, stimulates NADPH oxidase (NOX), producing cytosolic ROS ^42^. Additionally, mechanical stress from muscle contractions amplifies ROS production through the activation of mechanosensitive elements and increased mitochondrial respiration ^43^. Given the significant complexity of these cellular processes and the limited availability of data, we used a minimal linear model to account for PI3K/Akt, PKC, and stress-induced ROS production in the cytosol.

### Fusion-Fission Signaling Module

To model signaling pathways of mitochondrial fusion and fission in skeletal muscle during exercise, we employed a logic-based ODE modeling approach, which has been used extensively in modeling signaling pathways of cardiac and skeletal muscles ^33,37–39,45^. We chose this approach because it captures complex, large-scale signaling while requiring minimal time-course data, which is often unavailable in cardiac and skeletal muscle studies. The signaling network was constructed based on known interactions between key signaling molecules involved in muscle fusion and fission, such as calcium-calmodulin (Ca^2+^-CaM), p38 mitogen-activated protein kinases, mammalian target of rapamycin (mTOR), extracellular signal-regulated protein kinases 1 and 2 (ERK), AMP-activated protein kinase (AMPK), dynamin-related protein 1 (Drp1) and Mitofusin 1 and 2 ^46–50^. In total, the signaling model included 28 nodes representing different signaling species and 4 nodes representing the mediators of signaling and metabolic pathways (i.e., ATP/ADP, CPT1, ROS, and GLUT4) connected through 66 reactions. Each reaction uses a normalized Hill function with default parameters of weight (w), EC50, and Hill coefficient (n). Each signaling species has default parameters for initial level (Y0), maximum level (Ymax), and time constant (τ) ^36^. Details of the signaling model species, reactions, and parameters are provided in Supplementary File 1. Model inputs consisted of an Energy node, which was activated by Sprint, Endurance, and Resistance exercise regimens, and a sympathetic stimulation (Adrenaline) node, specifically activated by the Sprint exercise regimen. The model outputs were skeletal muscle fusion and fission.

### Logic-Based ODE Approach

A logic-based ODE (LDE) approach was utilized to transform the skeletal muscle fusion/fission signaling network into a system of ordinary differential equations, enabling the simulation of the signaling molecules dynamics during exercise ^36^. In the LDE approach, state variables *X* represent the normalized activity of each signaling species ranging between 0 and 1. Interactions between species are characterized as either activation *f_act_* or inhibition *f_inhip_*, represented by normalized Hill functions defined as follows:

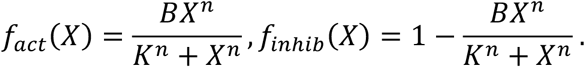

Here, 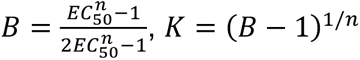. The LDE method uses continuous logic gates, AND (i.e., *f*(*x*)*f*(*y*) ) and OR (i.e., *f*(*x*) + *f*(*y*) − *f*(*x*)*f*(*y*) ) to model the interactions and crosstalk between signaling pathways in response to external stimuli. For example, PI3K activation can occur via either IGF1 or NRG1 receptor stimulation, which was modeled using an OR gate. However, muscle fusion requires the simultaneous activation of both MFN1 and MFN2, which was represented using an AND gate. The ODEs were generated automatically using Netflux, a software package designed for logic-based network modeling ^51^. Simulations were performed in MATLAB, with default parameter values based on previous studies ^36^. The model predictions were validated against experimental data on muscle signaling during exercise, ensuring that the model accurately captures the key dynamics of muscle fusion and fission.

### Sensitivity Analysis

To systematically evaluate the effect of individual reactions on mitochondrial fusion and fission dynamics, we implemented a global sensitivity analysis using the Morris elementary effects method ^52^. Morris’ method is well-suited for mechanistic models with high dimensionality or computational complexity, allowing for identification of parameters with the largest effect on system dynamics^53^. The weight parameters associated with each reaction in the network model were changed to examine their effect on the predicted levels of mitochondrial fusion and fission. To generate the input space for analysis, we adopted the Sampling for Uniformity (SU) approach. This sampling strategy was configured with 8 discrete levels for each input parameter, an oversampling pool of 300 candidate points, and 16 distinct trajectories to have a broad and uniform coverage of the parameter space ^54^. The sensitivity analysis was carried out using the EE sensitivity package in MATLAB developed by Khare et al. ^54^, which computed two key metrics for each reaction: the mean of the absolute values of the elementary effects (μ*) and their standard deviation (σ). The μ* value provides a measure of the overall influence of a reaction on selected outputs. By ranking the reactions in terms of μ*, we identified key reactions regulating mitochondrial dynamics (i.e., fusion and fission).

### Exercise input and conversion to MET

We considered three exercise regimens in this study: sprint, resistance, and endurance. To model their impacts on mitochondrial dynamics, we assumed that cellular energy level is influenced by exercise modalities, and the temporal changes in cellular energy follow the same temporal pattern as the exercise protocol described in Ref. ^3^. The basal cellular energy level was set to 0.2 for all three exercise regimens to simulate rest condition. For sprint exercise, the cellular energy level was elevated by 0.8 units above the basal level during the intervals [0, 0.5 min], [4.5, 5 min], and [9, 9.5 min]. For resistance exercise, the cellular energy level was elevated by 0.2 units above the basal level during the intervals [0, 0.5 min], [2.5, 3 min], [5, 5.5 min], [7.5, 8 min], [10, 10.5 min], and [12.5, 13 min]. For endurance exercise, the energy level increased by 0.12 during the interval [0 90 min]. These cellular energy levels were chosen to represent the different energy demands of the exercise regimens ^3^.

### Code availability

The mathematical model was simulated in SimBiology in MATLAB 2025b. The corresponding SimBiology project file is available in https://github.com/mkm1712/Muscle-Mitochondrial-Dynamics-in-Exercise.

## Results

### Classified Qualitative Validation of Fusion-Fission Signaling Network Model

To validate the accuracy of the developed systems biology model, we compared its predicted signaling activities to experimental data. The metabolic module had already been validated in a previous study ^31^. For the signaling model, the activity levels of key signaling molecules were compared to experimental data using a classified qualitative validation approach ^37^. Accordingly, the steady-state activity of signaling nodes following the exercise input (stimulated) was compared to their steady-state activity in the absence of the exercise input (control). Signaling nodes were categorized into three groups (“Model” row in Figure 2) based on relative changes in their activity compared to Threshold (i.e., 0.1% relative change): increased, decreased, and unchanged. Similarly, experimental data were classified into three categories, decreased, no change, or increased, based on their statistical significance when compared to controls reported in the literature (“Experiment” row in Figure 2). By conducting thorough manual review of existing studies, we identified experimental measurements for 22 nodes in the signaling network model, compiled from 18 distinct articles, which were independent of those used for model development. It is important to note that reports of exercise-induced signaling are not always consistent, with nodes such as PI3K/Akt, p38, mTOR, and AMPK showing opposing trends across studies. Such discrepancies likely reflect differences in experimental context, including cell state and methodological approach. The modular model correctly predicted the majority (80%) of observed signaling trends. This performance is comparable to other signaling network models and near the upper end of reported accuracies^33,37,39,45^. UQ studies showed this metric is robust to parameter uncertainty^55^.

**Figure 1:**
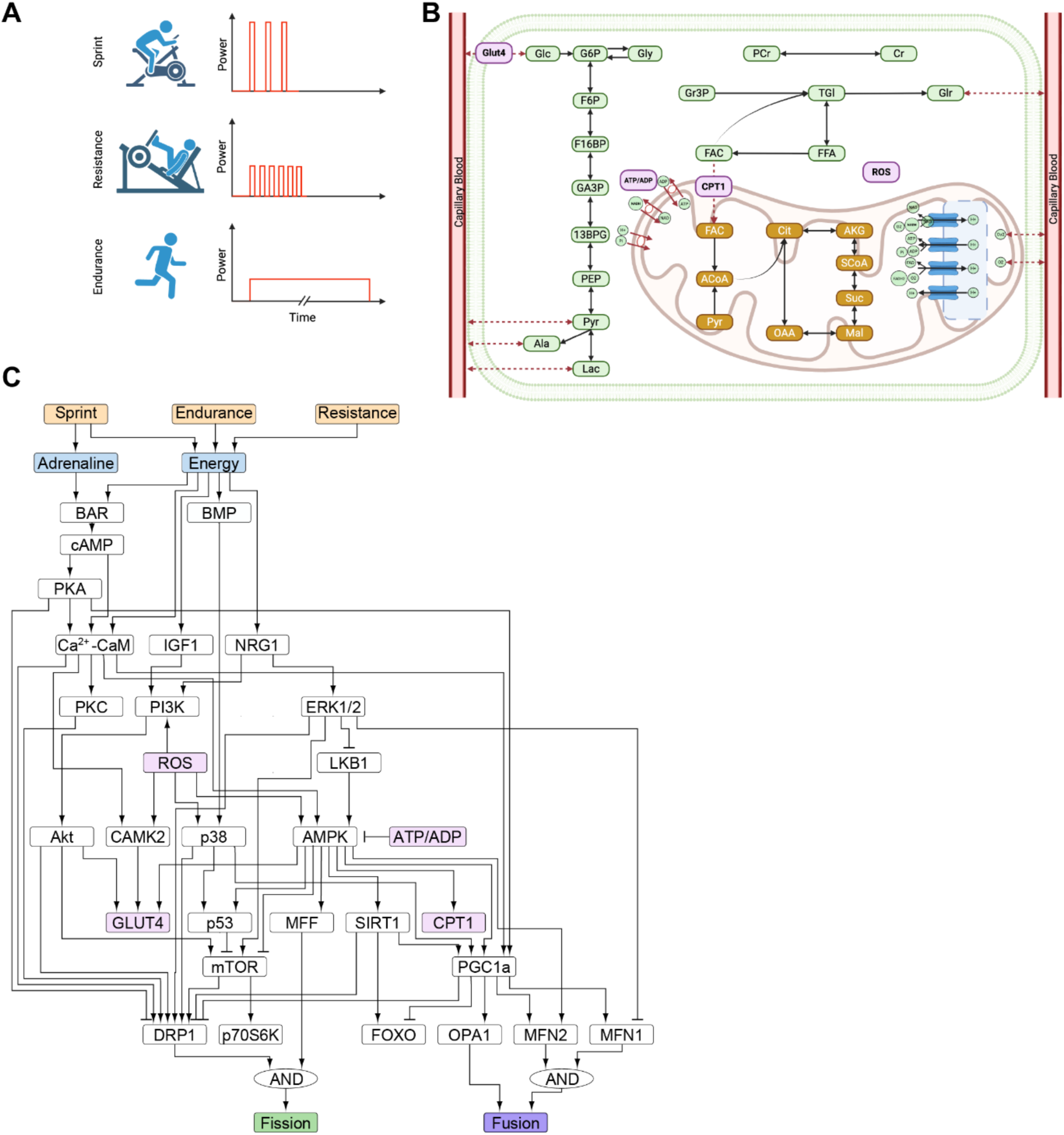
Systems biology model of mitochondrial dynamics. (A) Model time–power inputs illustrating the energetic load/stress of different exercise modalities. (B) Capillary– myofiber module that models GLUT4-mediated glucose uptake, glycolysis, PCr/Cr buffering, β-oxidation, TCA and ETC ATP generation, mitochondrial ROS generation, and metabolite exchange with blood. (C) Exercise signaling module linking adrenergic, Ca²⁺/CaM, IGF1/NRG1, PI3K-Akt, ERK-p38, AMPK, mTOR-SIRT1-PGC1α pathways to mitochondrial fission/fusion. Key mediators of signaling and metabolic modules (i.e., ATP/ADP, CPT1, ROS, GLUT4) highlighted in pink.

**Figure 2:**
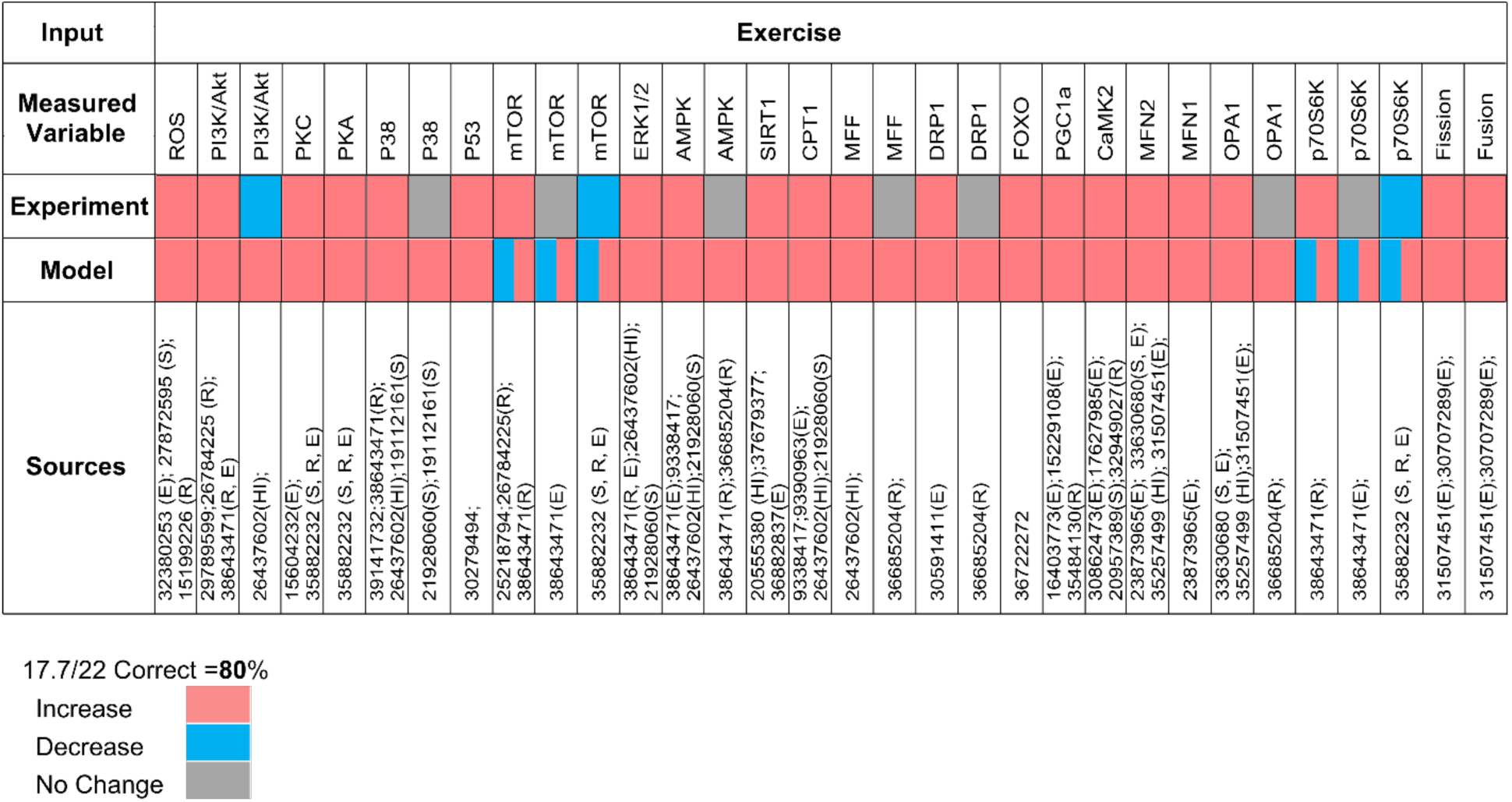
Classified Qualitative Validation of Fusion-Fission Signaling Network Model. Model predicted changes in fractional activation of signaling nodes are classified as increased (>0.1%, red), decreased (<–0.1%, blue), or unchanged (–0.1% to 0.1%, gray). Corresponding statistically significant experimental changes for the same molecules are shown for comparison, with supporting studies cited by their PMID number in Sources.

### Model predicts the dynamics of cytosolic species across exercise modalities

Having validated the modular model, we then investigated the normalized dynamics of cytosolic species under different exercise regimens. It is noteworthy that as all simulated responses are normalized and the magnitudes of absolute changes from pre-exercise level are not quantified. So, results should be interpreted qualitatively (e.g., cAMP activity is higher in sprint than endurance), not quantitatively. During exercise, sympathetic catecholamines activate β-adrenergic receptors in skeletal muscle, elevating cAMP–PKA signaling to mobilize fuel ^56–58^, while contraction-evoked Ca²⁺ transients engage Ca²⁺-calmodulin pathways ^59,60^, collectively increasing energetic demand. At the same time, depending on fiber type and exercise intensity, exercise could enhances glucose uptake and its hexokinase-mediated phosphorylation to glucose-6-phosphate, and accelerate anaerobic glycolysis, thereby increasing lactate ^61,62^. However, the model predicts distinct, modality-dependent cytosolic dynamics across exercise regimens. For cytosolic cAMP and Ca^2+^-CaM, the highest peak levels are observed during sprint exercise, followed by endurance exercise, with resistance exercise resulting in the lowest levels (Figure 3A-B). Cytosolic glucose and lactate show clear modality dependence: endurance evokes the largest and most sustained rises, sprint produces smaller, transient elevations, and resistance yields minimal changes. Peak amplitudes of glucose and lactose rank endurance > sprint > resistance (Figure 3C-E). Collectively, model predicts modality-specific dynamics for cytosolic species with maximum cAMP/Ca^2+^-CaM activity in sprint exercise, and highest cytosolic glucose/lactate level in endurance training.

**Figure 3:**
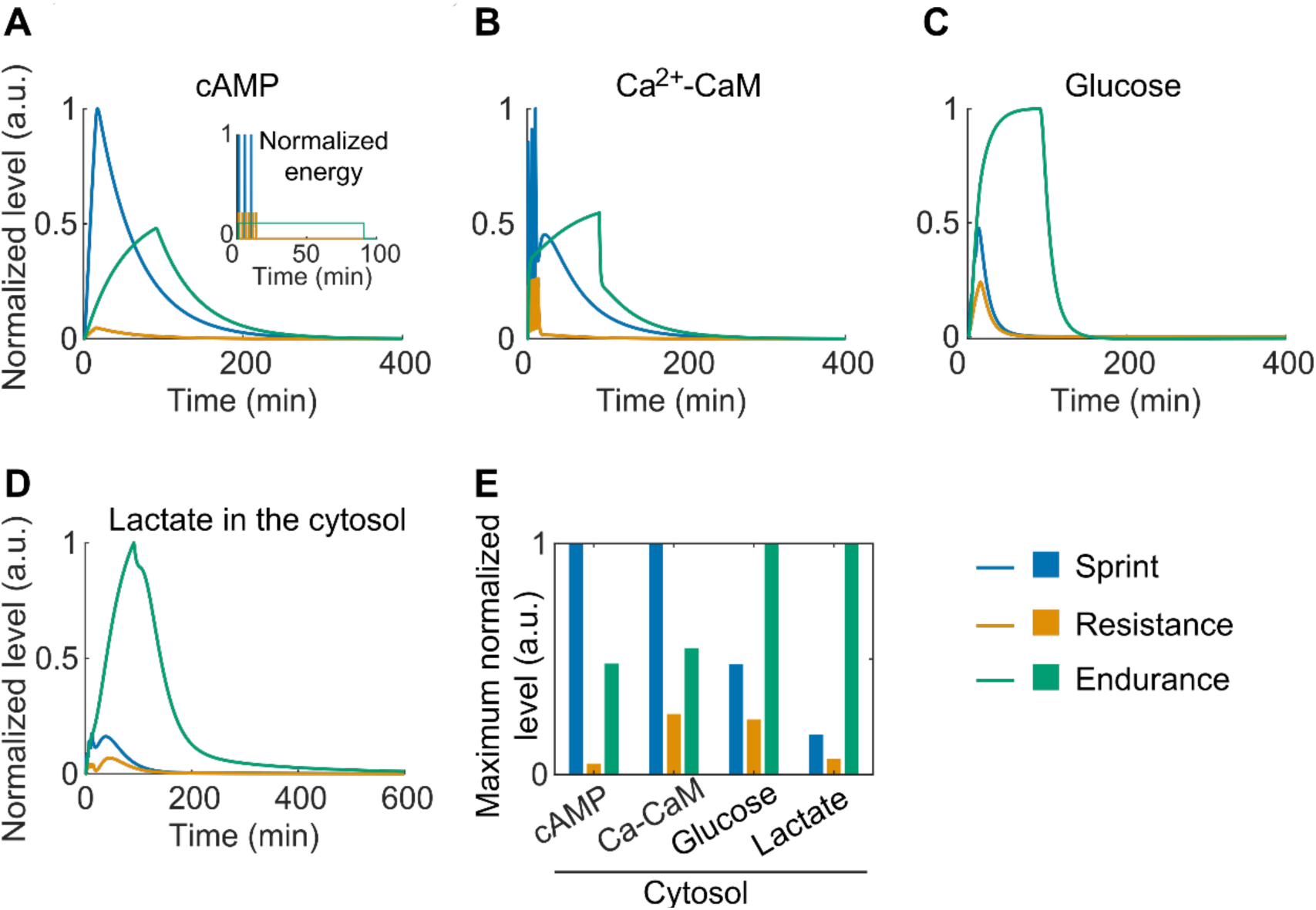
Predicted normalized dynamics of cytosolic species across exercise modalities. (A–D). Simulated dynamics of cytosolic cAMP (A), Ca^2+^-CaM (B), glucose (C), and lactate (D). In each panel, three different types of exercise regimens are applied: sprint (blue), endurance (orange), and resistance (green). The corresponding energy response is shown in the inset of panel (A). The normalized level is obtained by the fractional activation of species divided by the maximum changes in the fractional activation across three different types of exercise regimens. (E) Maximum normalized level for cytosolic cAMP, Ca^2+^-CaM, glucose, and lactate. The color indicates the exercise regimen: sprint (blue), endurance (orange), and resistance (green).

### Exercise-driven shifts in ROS, ATP/ADP, and AMPK–mTOR regulation

We then investigated the effects of exercise regimens on mediators of signaling and metabolism in skeletal muscle, focusing on cytosolic ROS, ATP, ADP, AMPK, and mTOR. Cytosolic ROS, a key metabolite positively regulated by PI3K, PKC, and NRG1, increases in response to exercise stimulation and relaxes during recovery (Figure 4A). Among exercise types, endurance training induces the highest ROS peak, followed by sprint and resistance training, consistent with greater electron-flux and oxidant production under prolonged load (Figure 4A, F). Cytosolic ATP and ADP levels also undergo significant changes during exercise. As exercise starts, ATP transiently decreases as it is hydrolyzed and consumed through various metabolic reactions to meet the energy demands of skeletal muscle, leading to a corresponding increase in ADP (Figure 4B-C). Notably, the dynamics of cytosolic ADP closely follow the energy profile, while ATP levels exhibit an inverse relationship to energy dynamics (Figure 4B-C). These energetic transients propagate to kinase control. AMPK exhibits a modality-dependent surge that is sharp and large with sprint, sustained with endurance, and minimal with resistance -- reflecting the predicted sensitivity of AMPK to changes in adenylate charge and glycogen state (Figure 4D,F). mTOR shows an early bump followed by suppression driven by AMPK/p53 inputs; this suppression is strongest and longest during endurance, weak and short-lived during sprint, and near-baseline with resistance (Figure 4E-F). mTOR activity rises after sprint and resistance exercise but declines following endurance. In our model, however, the recovery phase yields mTOR activity below the pre-exercise baseline for all regimens. In summary, our model predicts that endurance training elicits the largest, longest shifts in ROS and ADP, with sustained AMPK activation and strong mTOR suppression. Sprint exercise produces sharp, transient AMPK spikes, whereas resistance training yields minimal changes across these mediators.

**Figure 4:**
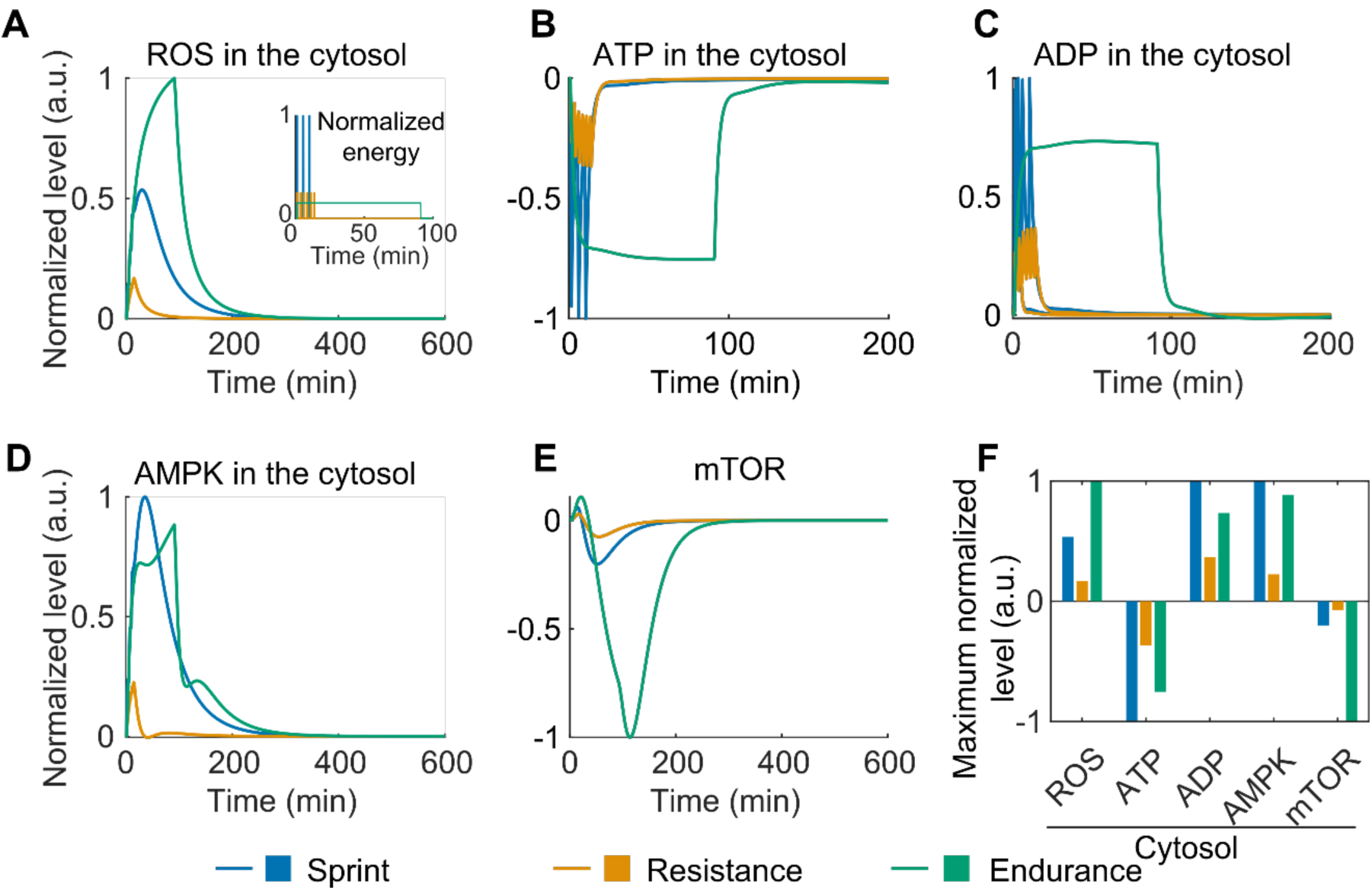
Predicted normalized dynamics of signaling and metabolism mediators across exercise modalities. (A-E) Simulated dynamics of cytosolic ROS levels (A), ATP levels (B), ADP levels (C), AMPK activity (D), and mTOR activity (E) in different exercise regimens. (F) Maximum normalized level for cytosolic ROS, ATP, ADP, AMPK, and mTOR. The color indicates the exercise regimen: sprint (blue), endurance (orange), and resistance (green).

### Regimen-specific control of mitochondrial ATP and the fission–fusion machinery

Next, we predicted how exercise modality shapes mitochondrial ATP flux and the fission-fusion machinery. Simulations predict a rapid rise in mitochondrial ATP production that tracks the imposed energetic load for all exercise modalities and relaxes with recovery (Figure 5A), yielding a transient increase in mitochondrial ATP that returns to baseline once demand subsides (Figure 5B). Endurance produces the largest and most sustained ATP-producing flux, sprint a sharp but brief burst, and resistance minimal change (Figure 5H). Upstream contraction and catecholamine signals (Ca²⁺-CaM, PKC, Akt, p38, ERK) drive a fast activation of DRP1, producing an early fission surge that diminishes after the stimulus (Figure 5C, F). In parallel, MFN1 and MFN2, activated via PGC-1α, ERK, and AMPK, accumulate more slowly, supporting a delayed fusion response that helps restore network connectivity and distribute the newly generated ATP (Figure 5D, E, G). Across readouts (mitochondrial ATP, DRP1/MFN activities, fission, fusion), peak amplitudes rank endurance > sprint > resistance (Figure 5H). Thus, the model predicts that modality-specific energetic load drives a staged remodeling sequence, an early DRP1-driven fission surge followed by MFN1/2-mediated re-fusion, with endurance producing the largest and most sustained mitochondrial reorganization.

**Figure 5:**
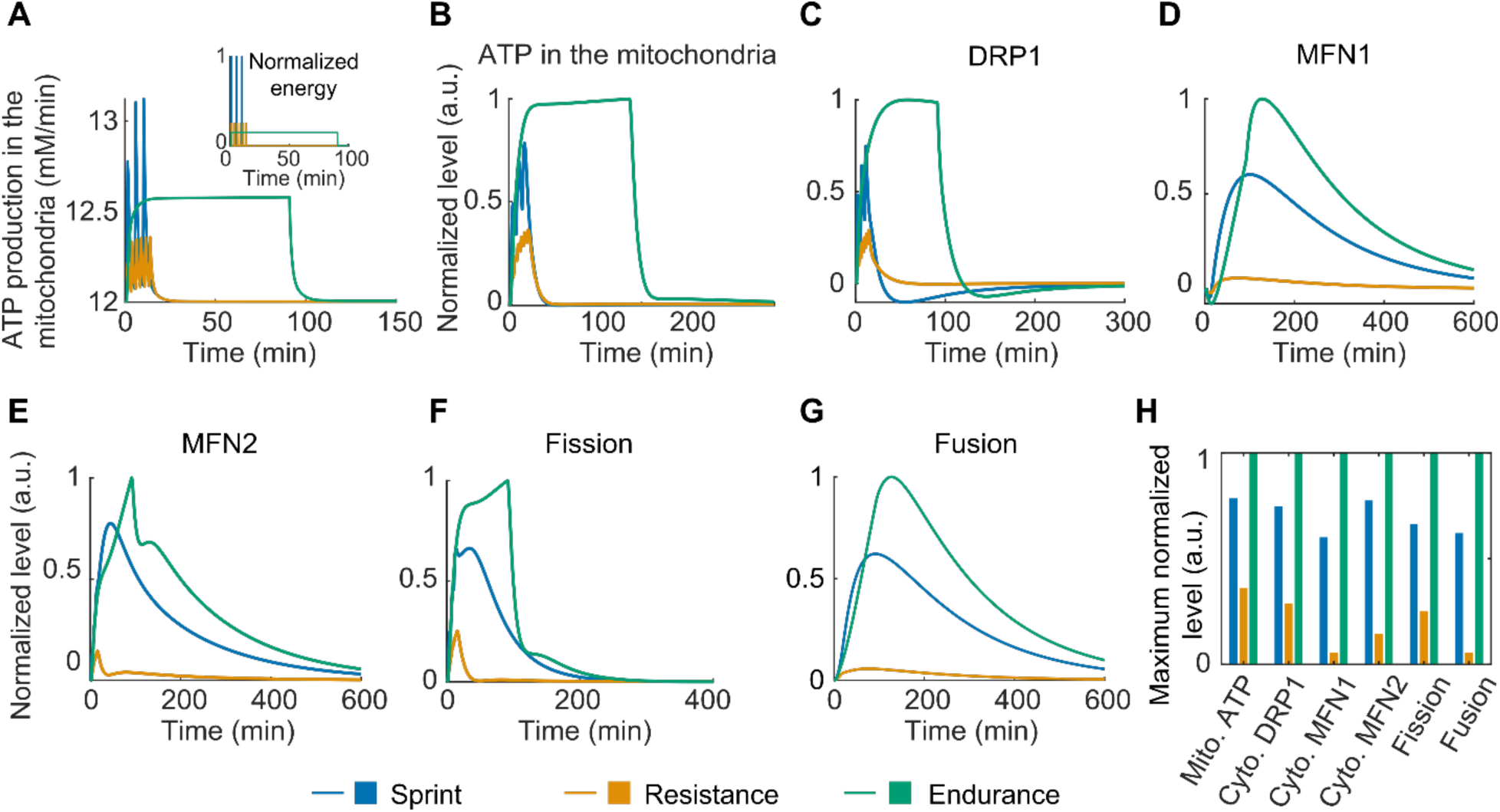
Predicted normalized dynamics of mitochondrial ATP and the fission– fusion machinery across exercise modalities. (A–G). Simulated dynamics of mitochondrial ATP production rate (A), mitochondrial ATP levels (B), cytosolic DRP1 activity (C), cytosolic MFN1 activity (D), cytosolic MFN2 activity (E), mitochondrial fission (F), and mitochondrial fusion (G) in different exercise regimens. (H) Maximum normalized level for mitochondrial ATP levels, cytosolic DRP1 activity, cytosolic MFN1 and MFN2 activities, and mitochondrial fission, and fusion.

### Sensitivity analysis identifies key control points for mitochondrial fusion and fission

To identify reactions with major influence on mitochondrial dynamics during exercise, we conducted a global sensitivity analysis using fusion and fission activities as readouts. This is a common approach to identify major signaling interactions regulating the cell response ^33,63^. Varying the weight parameter (w) of each reaction across the network revealed a compact set of high-leverage pathways (Figure 6). On the fusion arm, AMPK/PGC-1α signaling and the effector nodes MFN1/MFN2 exert the strongest positive control. On the fission arm, ROS input, AMPK-dependent signaling, and the DRP1–MFF axis dominate. Several mediators (e.g., Ca²⁺-CaM, ERK1/2, LKB1, and AMPK) influence both processes indicating shared upstream switches that can flip network polarity depending on energetic and calcium cues. Overall, the sensitivity analysis highlights AMPK as a central signaling hub and suggests that enhancing AMPK/PGC-1α/MFN1/2 biases skeletal muscle mitochondria toward fusion, whereas amplifying ROS/DRP1/MFF drives fission.

**Figure 6:**
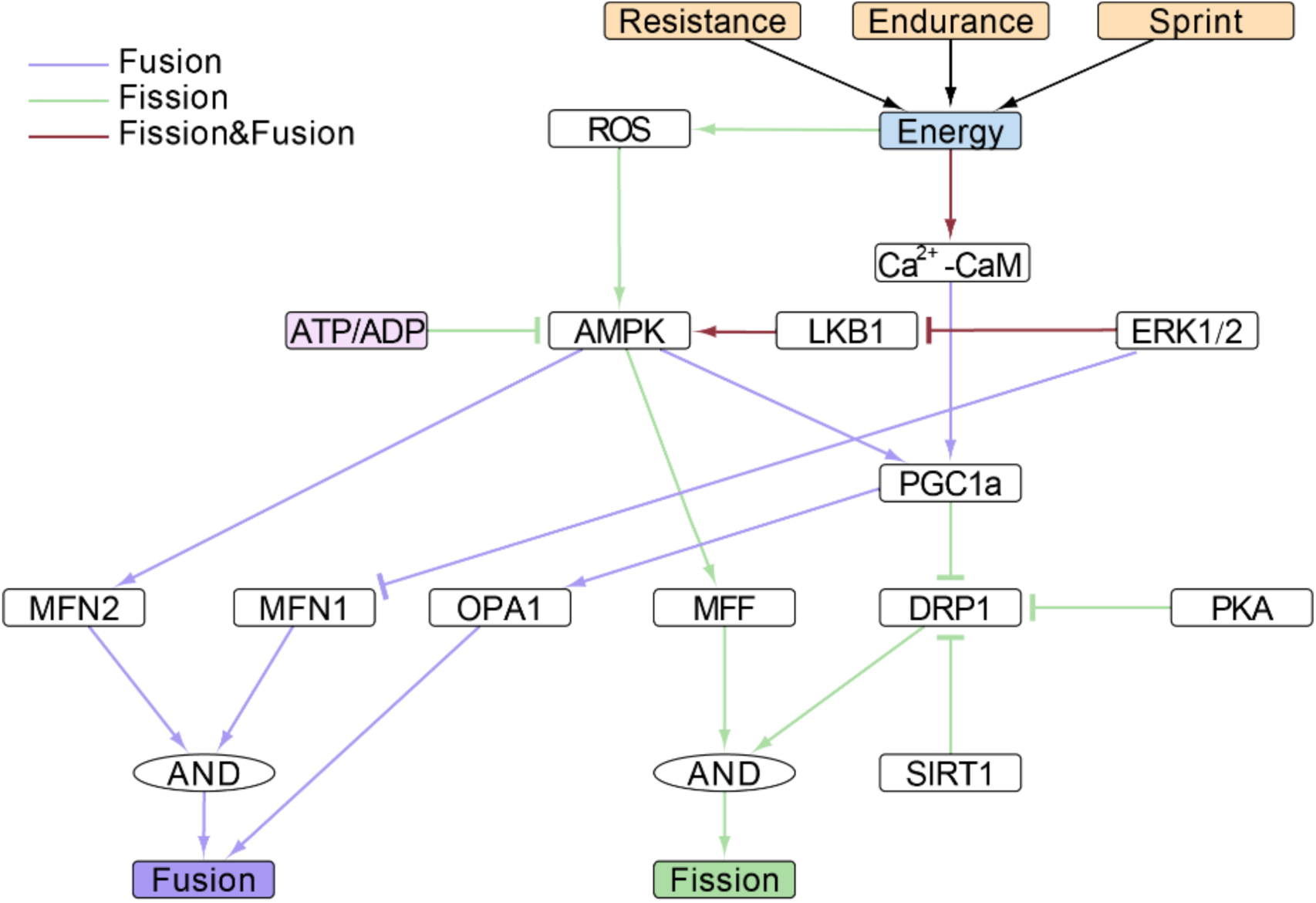
Sensitivity analysis identifies major reactions regulating mitochondrial fusion and fission. Morris sensitivity analysis identified major reactions that control mitochondrial fusion (purple), fission (green), or both processes (red). Exercise modalities (sprint, endurance, resistance) provide the energetic/Ca²⁺ inputs that drive signaling through Ca²⁺–calmodulin, ERK1/2, LKB1–AMPK, ATP/ADP, and ROS. AMPK and PGC-1α emerge as central integrators that transmit these cues to the mitochondrial machinery: fusion effectors MFN1/2 and OPA1, and fission effectors DRP1 and MFF. Logical “AND” gates indicate that both inputs are required. Arrows and bars denote activation and inhibition, respectively.

## Discussion

Here, we developed a modular computational framework that integrates blood–myofiber energetics, cytosolic signaling networks, and the mitochondrial fission–fusion machinery to explain why sprint, resistance, and endurance exercise elicit distinct yet overlapping mitochondrial adaptations in skeletal muscle. The simulations reproduce a stereotyped temporal sequence: an acute DRP1-dependent fission burst during the exercise bout, followed by MFN1/2- and OPA1-mediated re-fusion during recovery as energetic stress resolves and gene regulatory programs (e.g., PGC-1α–driven) mature. These results align with a view of exercise as a cyclical perturbation that transiently fragments the network to triage damage and later restores connectivity to distribute newly synthesized components ^64^.

### ROS-mediated control of mitochondrial dynamics in exercise

Predictions from our model indicate that reactive oxygen species (ROS) are early drivers of the shift toward mitochondrial fission, and their production rises with metabolic load and with catecholamine- and Ca²⁺-dependent activation of NOX2 and respiratory-chain electron flux ^65^. By examining whether antioxidant vitamin supplementation alters endurance-training adaptations in humans, researchers indicated that physiological ROS are required for normal adaptation ^66^. Similarly, high-dose vitamins C and E in a placebo-controlled trial impaired cellular markers of endurance adaptation, supporting that ROS signaling is beneficial during training rather than purely damaging ^67^. Results from the model also show that endurance exercise produces the largest and most sustained cytosolic ROS increase together with the strongest DRP1 activity and fission, in line with studies reporting exercise-induced increases in DRP1 expression or phosphorylation and a shift back toward fusion during recovery^68^. Mechanistically, AMPK links energy stress to mitochondrial fission by phosphorylating MFF, which recruits DRP1 and promotes fission early after exercise ^69^. By contrast, PKA phosphorylation of DRP1 at Ser637 restrains fission and favors fusion, so the final morphology reflects the balance between stress-activated kinases and cAMP–PKA signaling during and after contractions ^70^. In summary, ROS and energy stress act together to bias the network toward fission at exercise onset, with fusion re-emerging as redox pressure and catecholamine tone return toward baseline in recovery^68,70^.

### AMPK tunes the balance between mitochondrial fission and fusion during exercise

AMPK serves as the cell’s energy hub, linking rapid, activity-driven shifts in mitochondrial morphology to slower gene-regulated remodeling programs. It rises quickly during sprints and remains elevated with endurance exercise, translating energy stress into actionable signals in vivo. Specifically, AMPK activates ULK1 to promote autophagy and phosphorylates MFF to recruit DRP1, initiating mitochondrial fission^69,71^. Over longer time scales, AMPK together with p38 activates PGC-1α, which induces genes for fusion (MFN1, MFN2, OPA1) and oxidative machinery. In humans, both endurance training and HIIT rapidly increase nuclear PGC-1α and its mRNA, with fusion rising later after an initial fission phase ^14,72,73^. Exercise elevates MFN2 and this increase tracks with mitochondrial function, whereas reduced reliance on oxidative metabolism lowers MFN2, linking cellular energy status to fusion capacity ^74^. Genetic removal of MFN1/2 in adult mouse muscle impairs electron-transport chain function and diminishes exercise performance, demonstrating that a competent fusion program is essential for training benefits^75^. Taken together, AMPK-dominant signals initiate early fission for triage and mitophagy, followed by AMPK–p38–PGC-1α-driven fusion and biogenesis that rebuild a more connected, higher-quality mitochondrial network ^76^.

As AMPK sits at the crossroads of these pathways, drugs that raise or lower AMPK can change how exercise reshapes mitochondrial fission and fusion. The direction depends on timing, dose, and context (e.g., exercise modality and muscle state) ^76^. Direct AMPK activators such as A-769662, and 991 could mimic parts of the exercise signal in type II muscle fibers. They enhance glucose uptake and promote mitochondrial biogenesis. They can also drive fission-linked mitophagy via ULK1 and MFF ^69,71,77^. A key caveat is that A-769662 preferentially activates β1-containing AMPK complexes, which may not reproduce contraction-relevant γ-subunit signaling; thus, ex vivo findings may not translate to training adaptations ^77^. Prolonged AMPK activation also carries trade-offs. It can suppress mTOR–S6K signaling and alter ribosome biogenesis, blunting hypertrophy from resistance exercise and, in some cases, limiting gains in mitochondrial respiration even when mitochondrial “content” increases ^78,79^. In older adults, metformin, an indirect AMPK activator, may reduce muscle growth during structured training and attenuate improvements in cardiorespiratory fitness, indicating that layering AMPK activators onto exercise can be counterproductive depending on the objective ^78,79^. There are contexts, however, where activation is beneficial. For example, when mitochondrial quality is poor, as in certain dystrophy models, 991 or AICAR improved muscle homeostasis and regeneration, consistent with a setting in which enhanced mitophagy and fission-mediated triage are desirable ^80^. Overall, acute and potent AMPK activation is expected to amplify the early fission/mitophagy phase of an exercise bout, whereas chronic use around resistance training may dampen the later fusion/biogenesis phase via mTOR suppression; careful periodization and avoidance of indiscriminate AMPK agonism when hypertrophy is the priority are therefore warranted ^76^.

### Optimizing exercise for mitochondrial health

A key practical question is which training plan best supports mitochondrial health when we consider both fission and fusion. Our simulations indicate that workouts with higher total energetic level like endurance as well as sprint (high-intensity intervals) create the biggest short-lived AMPK–ROS signal and DRP1 activation, followed by strong fusion as PGC-1α–driven programs turn on. In contrast, low-load resistance work causes smaller redox and AMPK changes and milder effects on dynamics, even though it clearly benefits strength and ribosome biogenesis ^14,81,82^. Human studies show that both single and repeated interval sessions boost nuclear PGC-1α and other markers of mitochondrial biogenesis, and in some cases achieve gains similar to traditional continuous endurance training but in much less time ^14,81,82^. There is an active debate about what matters most: intensity or volume. Studies suggest volume mainly drives increases in mitochondrial content (e.g., citrate synthase), whereas intensity is a stronger driver of mitochondrial function (respiratory capacity)^83^. This view aligns with our finding that similar fission-fusion patterns can arise from different combinations of intensity and exercise duration. Protein data support a “fragment-then-fuse” pattern. MFN2 and OPA1 tend to rise with both moderate continuous training (MICT) and sprint/HIIT, while DRP1 phosphorylation spikes during exercise and returns toward baseline in recovery ^68,84^. Head-to-head studies further suggest that high-intensity aerobic sessions can produce larger, better-connected mitochondrial networks than resistance-only or mixed protocols over similar time frames ^85^. Therefore, a practical plan for mitochondrial health could be an endurance base plus one to two HIIT/SIT sessions per week to amplify the AMPK–ROS pulse and its downstream mitophagy/fusion program ^86^. It could be complemented with at least one resistance session to preserve myofibrillar quality and the translational capacity needed for mitochondrial protein renewal ^87^. In simple terms, longer low-to-moderate sessions are especially good for boosting mitochondrial content, whereas high-intensity intervals are especially good for improving mitochondrial function; combining them likely maximizes both while avoiding overreliance on a single pathway ^81,83^. Mechanistically, acute endurance exercise raises PGC-1α transcription and nuclear levels in human muscle, with promoter demethylation strengthening this response and providing the substrate for the delayed fusion/biogenesis phase ^20,72,73^. By contrast, antioxidant supplementation that blunts the exercise ROS signal dampens these transcriptional programs, supporting the need for a sufficient redox “pulse” to help the system move from fission during exercise to fusion in recovery ^67,88^.

### Limitations and future directions

Our model offers a modular way to connect exercise signals to mitochondrial fission and fusion, capturing qualitative trends and pathway crosstalk without needing many hard-to-measure kinetic constants. As such, it is well suited for semi-quantitative predictions and hypothesis generation. However, there are some methodological limitations that may restrict the generalizability of the results. The logic-based differential equation approach cannot provide absolute molecule concentrations or fully capture gradual, time-dependent enzyme kinetics, so outputs should be interpreted as semi-quantitative. Also, the current ROS module compresses multiple oxidant sources into a single minimal representation. Experiments indicate that NOX2 is a major source of cytosolic ROS during endurance-type exercise and that mitochondrial ROS can even drop briefly, suggesting a need for compartment-specific redox modules that explicitly include NOX2/NOX4 and respiratory-chain leak ^65^. In this model, the muscle is treated as one uniform compartment. Whereas real muscle differs by fiber type, recruitment pattern, and mitochondrial subpopulations. Thus, adding fiber-resolved energetics and spatially explicit mitochondria would better reflect this heterogeneity ^88^. While the model includes DRP1, MFN1/2, and OPA1 dynamics but does not yet model OPA1 processing, cristae remodeling, or the MICOS complex. These features influence how fusion translates into respiratory efficiency, so incorporating these processes could enhance model predictions for how endurance versus interval training remodel the inner membrane. Finally, we have not yet linked mitochondrial dynamics to mitophagy flux beyond ULK1 targeting. As fission helps isolate damaged segments and mitophagy is required for many training benefits, adding a quantitatively constrained turnover module enhances model predictions ^89^. Despite these constraints, the model’s strengths including its network-based structure, modularity, and the ability to integrate diverse exercise signals, make it a practical framework for testing mechanism-based interventions and for prioritizing experiments. By adding compartmental redox modules, spatial/fiber-type structure, inner-membrane remodeling, and explicit mitophagy flux in future studies, the same framework can deliver more precise predictions.

## Conclusions

In summary, the present model supports a unifying principle: exercise reshapes mitochondria in phases. An AMPK/ROS-driven fission burst isolates damage and triggers mitophagy. Then AMPK-p38-PGC-1α drives refusion and biogenesis to rebuild the network. Exercise intensity and volume set the size and timing. From a prescription standpoint, we posit that a blended program with an endurance base, periodic HIIT to intensify the early signal, and regular resistance training to maintain translational capacity will best sustain mitochondrial health through coordinated fission–fusion–turnover. We predicted that indiscriminate chronic AMPK pharmacology can, in some populations, attenuate training adaptations and should be periodized or avoided when hypertrophy or maximal respiratory gains are desired. Finally, the model could provide a scaffold for testing targeted hypotheses (e.g., NOX2 inhibition, β-adrenergic blockade, or fiber-type– specific recruitment) against the still-growing experimental studies on mitochondrial dynamics in human muscle.

## Acknowledgments

This work was supported by the Wu Tsai Human Performance Alliance at the University of California, San Diego

## Supplement

**Figure S1:**
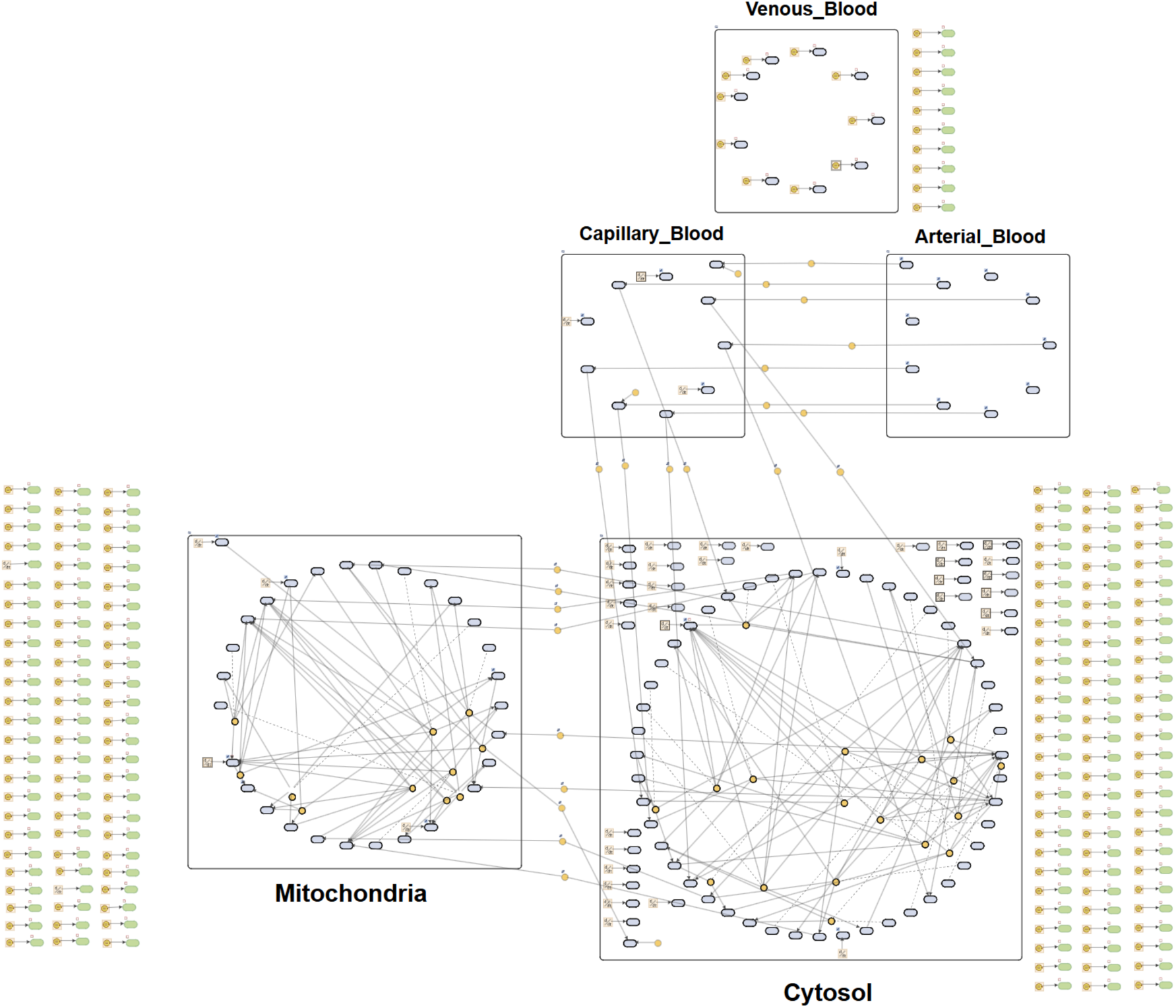
Multi-compartment skeletal-muscle signaling–metabolism model in SimBiology. The model topology links blood (arterial, capillary, venous), cytosol, and mitochondria. Boxes denote compartments; blue nodes represent dynamic species; arrows indicate reactions, transport, and regulatory influences (solid = flux/reaction; dashed = regulation). Circulating substrates and oxygen enter from blood, feed cytosolic pathways, and couple to mitochondrial energetics.

**Supplementary File 1: List of Species, initial conditions, reactions and parameters of the logic-based ODE signaling model**

